# Taxonomic practice, creativity, and fashion: What’s in a spider name?

**DOI:** 10.1101/2022.02.06.479275

**Authors:** Stefano Mammola, Nathan Viel, Dylan Amiar, Atishya Mani, Christophe Hervé, Stephen B. Heard, Diego Fontaneto, Julien Pétillon

## Abstract

There’s a secret pleasure in naming new species. Besides traditional etymologies recalling the sampling locality, habitat, or morphology of the species, etymologies may be tributes to some meaningful person (for example, the species’ collector, the author’s husband or wife, or a celebrity), pop culture references, and even exercises of enigmatography. The possibility of choosing witty or even playful names for new species departs from the otherwise impersonal and old-fashioned writing style that’s common in taxonomic papers; but, how has the descriptor’s choice for specific etymologies changed over the 300+ years since the introduction of the Linnaean binomial system of nomenclature? Using an unprecedented dataset of 48,464 etymologies (all valid species and subspecies of spiders described between 1757 and May 2020), we tested the hypothesis that species names given by taxonomists are deeply influenced by their cultural background. In particular, we asked whether naming practices change through space (continent in which the species was found) or have changed through time (year of description). We observed spatial and temporal differences in the way taxonomists name new species. In absolute terms, etymologies referring to morphology were the most frequently used. In relative terms, however, references to morphology peaked in 1850–1900 and then began to decline, with a parallel increase in etymologies dedicated to people and geography. Currently, these are the most widely used, with ∼38% of all etymologies of spider species described in the last ten years referring to geography, ∼25% to people, and ∼25% to morphology. Interestingly, there has been a dramatic increase in etymologies referring to pop culture and other cultural aspects in the last two decades, especially in Europe and the Americas. While such fashionable names often carry little or no biological information regarding the species itself, they help give visibility to the science of taxonomy, a discipline currently facing a profound crisis within academia. Taxonomy is among the most unchanged disciplines across the last centuries in terms of background, tools, rules, and writing style; but our analysis suggests that taxonomists remain deeply influenced by their living time and space.

## INTRODUCTION

In a quest to make some sense of the beautiful enigma often referred to as ‘reality’, humans have always attempted to classify natural objects and phenomena into rigid schemes and categories. The periodic table of elements by Mendeleev (Scerri, 2019), the categorization of stars according to their luminosity (Langer & Kudritzki, 2014), and the schematization of human behaviors into personality traits (Goldberg, 1993) are just a few, cherry-picked examples illustrating how the impulse to classify is inherent to any human attempt to understand the natural world (Doolittle, 1999). While some of these classifications may have more grounding than others in underlying reality, they reflect the fact that—as Richard Dawkins nicely put it—we are all subject to the “*tyranny of the discontinuous mind*”.

Among the most successful and long-lasting scientific classification schemes is the Linnaean binomial system of nomenclature (Linnaeus, 1758), proposed by Carl Linnaeus (1707–1778) three centuries ago and still in use to name living and fossil organisms. A species is unambiguously defined *via* the juxtaposition of two “Latin” names: the genus and the species epithet. Notwithstanding philosophical discussions around the biological meaning of these categories (Lehman, 1967; Hey, 2001; Slater, 2016), the heuristic of naming species has aided generations of scientists in formulating and testing hypotheses in disciplines as diverse as ecology and evolution (Pante *et al*., 2015; Fišer, Robinson, & Malard, 2018), agronomy (Malézieux *et al*., 2009), medicine (Yang *et al*., 2020), and conservation biology (Reydon, 2019).

Beyond the theoretical and practical usefulness of establishing unambiguous names to refer to species (Vink, Paquin, & Cruickshank, 2012; Zeppelini *et al*., 2021), the use of Linnaean nomenclature comes with an additional benefit for taxonomists: they get to choose species names. Besides a few basic grammatical rules regulating the derivation of etymologies (Vendetti & Garland, 2019), there are virtually no limits to authors’ creativity (Lalchhandama, 2014; Jozwiak, Rewicz, & Pabis, 2015; Heard, 2020). Names may describe species traits [the (blue) cerulean warbler, *Setophaga cerulea* (Wilson)]. They may refer to a species’ distribution (the Hawai’ian lobelioid tree *Cyanea heluensis* Oppenheimer, known only from the slopes of Mt. Helu; Oppenheimer, 2020) or habitat [the rock- and reef-inhabiting sergeant-major fish, *Abudefduf saxatilis* (Linnaeus)]. More creatively, they may refer to people (Kate Winslet’s beetle, *Agra katewinsletae* Erwin; or Margaret Collins’ ant, *Strumigenys collinsae* Booher), or simply represent wordplay (the tiny frog *Mini mum* Scherz et al. or the braconid wasp *Heerz lukenatcha* Marsh). The possibility of choosing witty or even playful names contrasts with the usually serious and impersonal writing style of traditional scientific papers (Heard, 2014). This is anecdotally confirmed by listening to a few episodes of the podcast ‘NewSpecies’ (@PodcastSpecies). The host, Dr. Brian Patrick, invites taxonomists to talk about newly described species; toward the end of each episode, he always poses a key question: “Why this name?” The different answers to this question by the invited speakers, and their enthusiasm in explaining their etymological choices, leave no doubt that there is a distinct pleasure in naming new organisms.

Here, we delve into authors’ choices of etymologies more systematically, taking a long view of the process of naming species to understand how authors’ choices vary in space and have changed over time. As considering the whole diversity of species across the Tree of Life would be hardly possible, we focused on spiders (Arachnida: Araneae). Spiders are a perfect test case for our etymological endeavor, because:

i. spider taxonomy is constantly curated and updated online in a central repository, the World Spider Catalog (2022);
ii. thanks to a unique online collaborative effort (Nentwig, Gloor, & Kropf, 2015), all literature on spiders is freely available, making it possible to easily inspect species descriptions to ascertain an original etymology (when reported);
iii. the geographical ubiquitousness of spiders (Piel, 2018), the great diversity of species (currently almost 50,000 valid taxa; World Spider Catalog, 2022), their diverse body forms, adaptations, and ecological strategies (Cardoso *et al*., 2011; Foelix, 2011; Mammola *et al*., 2017), as well as the existence of a broad community of academics and amateurs studying them (Platnick & Raven, 2013; Mammola *et al*., 2017; Jäger *et al*., 2021), all should lead to an ample variety of etymological roots for spider names.

Based on an unprecedented dataset of 48,464 unique taxonomic entities, we modeled the proportional use of different etymology types over time and across continents. Stemming from the fact that taxonomy is an old and conservative science still largely based on the use of Latin in naming, our null hypothesis was that there should be no significant variations in the way taxonomists have assigned etymologies since the Linnaean system was introduced. Conversely, if taxonomists are prone to be influenced by their cultural background, we would expect differences in etymological categories across periods and continents.

## MATERIAL & METHODS

### Backbone taxonomy

On 02 June 2020, we downloaded the backbone taxonomy of the World Spider Catalog (2020), consisting at the time of 48,464 unique taxonomic entities (47,956 species and 508 subspecies of spiders). The temporal span of the database goes back to 1757 when Carl Alexander Clerck (1709–1865) named 67 spiders using binomial nomenclature in his book *Svenska Spindlar* (Clerck, 1757). Interestingly, Clerck’s book precedes the publication of the 10^th^ edition of *Systema Naturae* (Linnaeus, 1758), considered the start of nomenclature, by one year. Therefore, in order to consider Clerck’s spider descriptions valid under the system of zoological nomenclature, *Svenska Spindlar* is deemed to be published on 1 January 1758 (International Code of Zoological Nomenclature; Article 3.1).

### Classification of etymologies

We classified species etymologies into six broad categories: those referring to i) morphology, ii) ecology & behavior, iii) geography, iv) people, v) modern & past culture, and vii) other. Note that a single etymology may have multiple meanings and therefore belong to multiple categories. Within each category, we further broke down the meaning of species names into subcategories (see details in the sections below). In all analyses, however, we used the six main categories above as grouping units, both for ease of discussion and because the distinction between subcategories was sometimes blurry (see section “**Cross-validation of inferred etymologies**”).

To assign etymologies, we first and foremost checked original descriptions; secondarily, we inferred etymologies based on our knowledge of Greek and Latin and by consulting previous etymological works (Bonnet, 1961; Nilsson, 2010) and internet resources (e.g., Wikipedia, Wikispecies). When we consulted an original description, we looked for the specific section describing the etymology—typically under headings such as “Etymology” or “*Derivatio nominis*”. If such a section was missing, we checked the paper’s text for hints to the meaning of the species epithet. Thanks to our knowledge of different languages, the help of collaborators (see **Acknowledgements**), and through the use of Google Translate, we were able to inspect papers written in Chinese, English, French, German, Italian, Japanese, Latin, Portuguese, Russian, Spanish, and Swedish.

For each species, we indicated in the database whether we based our classification of the etymology on the original description (‘Original’; n = 22,416), whether we consulted the original description but we had to infer the etymology because it was not reported (‘Original + Inferred’, n = 1,018), or whether we inferred it without consulting the original description (‘Inferred’, n = 23,890). We left blank those etymologies we couldn’t infer (‘Original + Unknown’; n = 1,140) and excluded these observations from analyses.

#### Etymologies referring to “Morphology”

We used this category when an etymology referred to the size of the spider (subcategory: ‘size’), the shape of the body or some body part (subcategory: ‘shape’), or the general aesthetic/appearance of the species (subcategory: ‘color’). Whenever the name indicated the similarity to another species (usually names ending in *-oides* or beginning with the prefix *sub-*), we scored it as size = 1, shape = 1, and color = 1 unless a morphological feature was specified (e.g., “… *similar in the shape of the cymbium*”).

#### Etymologies referring to “Ecology & Behavior”

We used this category when a species name referred to the ecology and habitat of the spider (subcategory: ‘ecology’) or some behavioral adaptation (subcategory: ‘behavior’).

#### Etymologies referring to “Geography”

We used this category when an etymology referred to the distribution of the species, including those names referring to the type locality.

#### Etymologies dedicated to “People”

We used this category when the etymology was dedicated to a scientist (subcategory: ‘scientists’) or other people (subcategory: ‘otherPeople’). Whenever the species was dedicated to the collector of the species, we always assigned it to ‘scientists’; this would include amateur collectors, who are being recognized for their contribution to the scientific process. We assigned the species to ‘otherPeople’ when the identity was not specified in the original description but it was possible to infer the species was dedicated to a person based on indirect evidence. For example, in old descriptions the species name was capitalized when referring to a person’s name or surname. When a species name was dedicated to a fictional person, we classified it only in the category ‘Modern & Past Culture’.

#### Etymologies referring to “Modern & Past Culture”

We used this category when the etymology referred to contemporary culture (subcategory: ‘modernCulture’) or past culture (subcategory: ‘pastCulture’). These may include references to mythology, local tribes, pop culture, music bands, and so on. We applied the concept of modern or past culture relative to the species descriptor. For example, what we considered ‘modernCulture’ for Eugène Simon (1848–1924) would be ‘pastCulture’ for Norman Platnick (1951–2020).

#### “Other” etymologies

We used this category when the etymology didn’t fit any of the previous categories. These include names that are puns or arbitrary combinations of letters, that refer to an anecdote related to the collection of the species, and many others.

### Cross-validation of inferred etymologies

In order to assess the internal consistency in etymology assignments among authors, as well as the validity of inferred etymologies (which represented 49% of the dataset) as compared to original etymologies, we cross-validated the dataset. We based cross-validation on 400 randomly picked inferred etymologies (∼1% of the total dataset). For each etymology, we checked the original description and scored whether our inference: i) matched, i.e., there was a clue in the text that the inferred etymology fitted the given name, like in *Nemosinga atra* Caporiacco, for which the abdomen is described as “completely black” in the original paper (*ater, atra -um* meaning “black, dark-colored”); (ii) didn’t match, i.e., there was evidence for another explanation to the given name in the original description; or (iii) neither matched nor mismatched, i.e. there was no indication to support the inferred etymology in the original paper, but nothing invalidated the deduction.

Of the 400 cross-validated etymologies, 57.75% matched with the original description, 16.75% did not match with the original description, and 25.50% were indeed not reported. Percentages of mismatch decreased to below 3% when the agreement was based on the match of a broad etymological category—in other words, in almost all cases, confusion was between subcategories (e.g., ‘size’ versus ‘shape’) and so was not influential for our analyses. In a second step, we also evaluated between-author congruence using three pairs of comparisons (DF, JP, and SM), each of 40 etymologies. The percentage of agreement was high, ranging from 86.67% match for subcategories to 100% match for category. Given these cross-validation results, we assumed the database to contain only trivial errors.

### Statistical analyses

We carried out all analyses in R version 4.0.3 (R Core Team, 2021). We used the package “ggplot2” version 3.3.4 (Wickham, 2016) for visualizations.

#### Assignment of species geographic distribution

We used the distribution column in our backbone taxonomy to classify species distributions at the continental level. Distribution strings for species in the World Spider Catalog are not standardized: they can mention single countries (e.g., “Namibia”), multiple countries (e.g., “Italy and Spain”), continents (e.g., “Oceania”), ranges (e.g., “from Europe to Korea”), and other geographical units (e.g., “Pyrenees”, “unknown”). We used the *wscmap* dataset from the R package ‘arakno’ version 1.1.1 (Cardoso & Pekar, 2022) to convert each distribution string into a list of corresponding 3-letter ISO country codes. For example, we translated “North America” to the list “CAN, MEX, USA,” and “Egypt to Yemen” to “EGY, SAU, YEM”. Next, we used the R package ‘countrycode’ version 1.3.0 (Arel-Bundock, Enevoldsen, & Yetman, 2018) to convert these lists of ISO codes into presence/absence values per continent (‘Asia’, ‘Europe’, ‘Africa’, ‘Oceania’, and ‘Americas’). We manually checked the final output, filling unmatched distributions and fixing errors.

#### Hypothesis testing

We tested our hypotheses using regression models. In all these analyses, we followed general recommendations by Zuur & Ieno (2016) for data exploration, model fit, and model validation. Given the large sample size, we used a conservative approach in the identification of significance, setting an alpha level for significance at 0.001 instead of the usually accepted 0.05 and always reporting effect size (Benjamin *et al*., 2018; Muff *et al*., 2021). In all analyses testing temporal patterns, we excluded the year 2020 because we had only partial data up to June.

To ask whether the way taxonomists choose etymologies has changed over time, we first calculated the annual proportion of etymologies for each etymology type (a factor variable with six levels, as described in the section “**Classification of etymologies**”). We performed a generalized additive model with the proportion of etymologies as a response variable, testing the effect of a smooth function of year of publication (a discrete variable from 1757 to 2019) in interaction with the etymology category. We used additive models in order to account for possible non-linear trends of the sampling period. We ran the model with the R package ‘mgcv’ version 1.8-35 (Wood, 2004) assuming a binomial distribution, a logit link function, and a thin plate regression spline. We estimated the optimal amount of smoothing through generalized cross-validation. As the initial model was overdispersed (dispersion ratio = 1.91), we switched to a quasi-binomial distribution, a quasi-distribution that includes an extra parameter (*ϕ*) to describe additional variance in the data that cannot be explained by a binomial distribution alone. The model sample size was 1,386 observations.

Next, we asked whether the temporal trends identified in the previous analysis changed across continents. We constructed a second model with the proportion of etymologies as a response variable, testing the effect of a smooth function of year of publication in interaction with the continent in which the species occurs (a categorical variable with five levels, see section “**Species geographic distribution**”). In this case, we stuck to a binomial distribution because overdispersion was minimal (dispersion ratio = 1.34). The model sample size was 6,378 observations.

Finally, we tested whether taxonomists are more likely to choose a given etymology category over others across continents. We constructed six generalized linear models, one for each etymology type, testing the effect of the factor continent on the use of a given etymology type. As the response variables were binary variables (0–1 discrete), we specified a Bernoulli distribution and a complementary log-log link function (clog-log), as recommended in Zuur et al. (2009) for datasets with unbalanced sets of zeros and ones. To avoid confounding effects of including species that occur across multiple continents, we used for the analysis the subset of species occurring in a single continent, resulting in a total sample size of 44,655 observations.

## RESULTS

### Spider etymologies by the numbers

We classified the meaning of 47,325 spider species and subspecies names. Of the classified names, most etymologies referred to spiders’ morphology (41%, n = 20,702) or geographic distribution (27%, n = 13,880), or were dedicated to people (19%, n = 9,881) (Figure 1A). The most frequently used names across the database also mostly belonged to these three categories (Figure S1). The least used etymologies belonged to the category Other (4%, n = 2,128). Over 90% of names attributed to species had a single meaning (i.e., we classified them into a single subcategory). There were, however, names with multiple meanings— typically two, although there were etymologies with up to four meanings. The average length of species epithets was 8 letters (Box 1).

**Figure 1.**
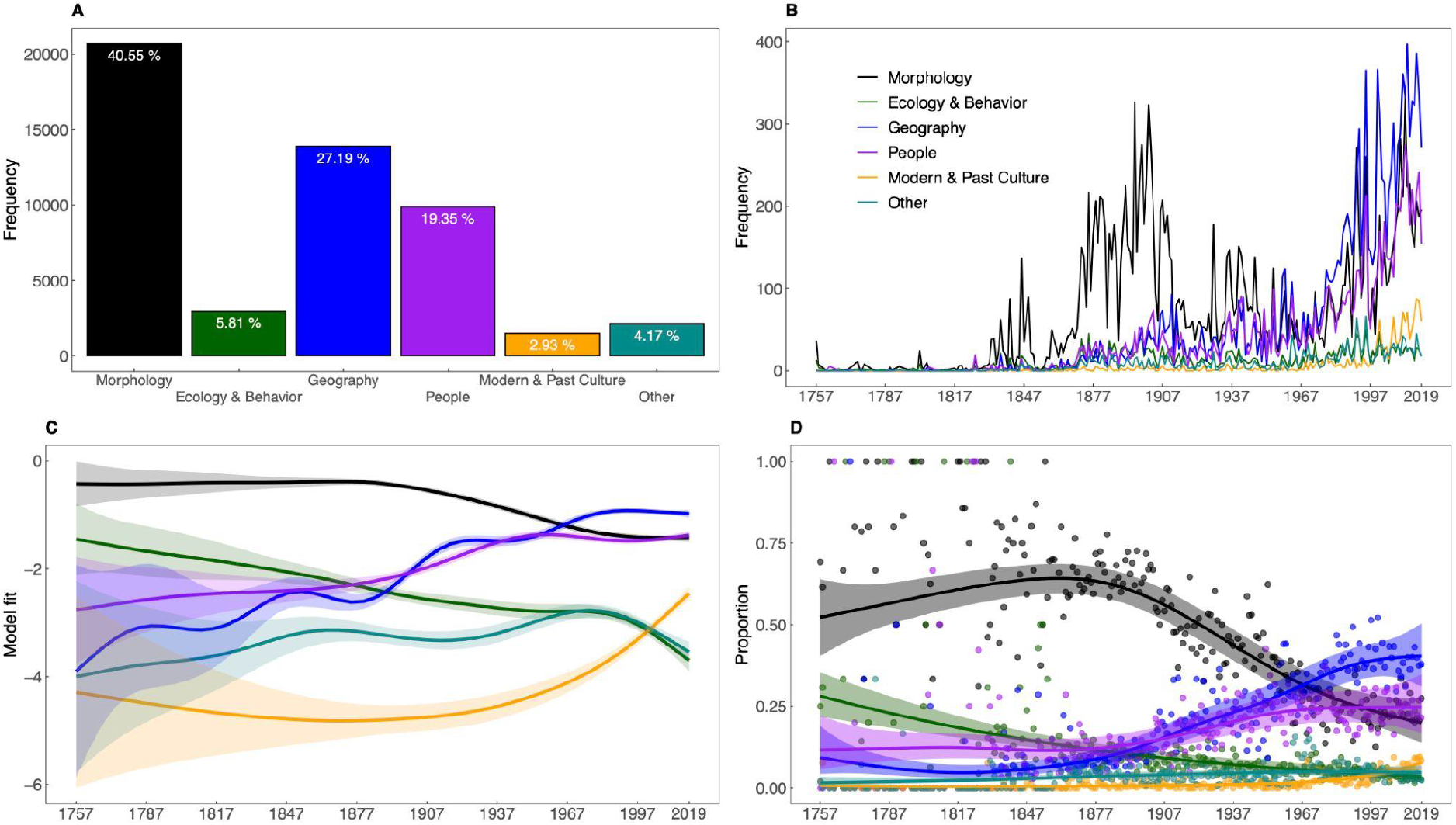
Temporal variations in the way taxonomists assign spiders etymologies. **A)** Breakdown of the total number of etymologies by category. **B**) Temporal variations in the number of etymologies (annual sum by etymology type) between 1757 and 2019. **C**) Plot of smooth terms for the temporal variations in the relative proportion of different types of etymologies, according to a generalized additive model. **D**) Temporal variations in the proportional use of etymologies. In **C** and **D**, colored lines and shaded surfaces are, respectively, the predicted trend and 95% confidence interval for each level of the factor type, according to a generalized additive model. Exact model estimates are given in Table S1.

### Temporal patterns

Are we naming species differently today than in the past? Looking at absolute numbers, there were some striking temporal differences in the way taxonomists name species (Figure 1B). This was confirmed by modeling the relative proportion of etymologies over time (Figure 1C), whereby we found a significant interaction between the year of the description and the type of etymology (R^2^ = 0.942; estimated parameters are in Table S1). In the first 150 years from the introduction of the Linnaean system of nomenclature, the most frequently used etymologies referred to morphology and ecology. The relative use of morphological etymologies peaked in 1850–1900 and then started to decline (Figure 1D) along with a parallel increase in etymologies dedicated to people and geographic distributions. Etymologies referring to geography quickly rose, overtaking morphological etymologies around 1950, and currently are the most widely used (∼38% of all etymologies of species described in the last ten years referred to geography, ∼25% to people, and ∼25% to morphology). Etymologies referring to modern and past culture began to rise in use after the year 2000. Although such etymologies are not widely used in absolute terms, their relative use has increased dramatically in the last two decades (especially those referring to pop culture).

A second model testing the interaction of etymology x continent revealed that the temporal trends differed significantly across regions, especially with respect to Oceania (Figure S2). The general direction of temporal trends across regions was, however, qualitatively similar and thus, we presented the full model in the main text (Figure 1D).

### Spatial patterns

While the proportions of names from the six etymological categories were quite consistent across continents (Figure 2), we found some notable spatial patterns in the way taxonomists assign species names (model estimates in Table S2–S7). Spiders from Europe are less likely to be named in reference to morphology, but more likely to bear names dedicated to some person, compared to spiders from other continents. Asian species were less likely to be named in reference to ecology & behavior than in other continents, whereas African species were less likely to be named in reference to geography. The probability of a species name being a reference to modern & past culture was higher in Europe and North America than in other continents (Figure 2).

**Figure 2.**
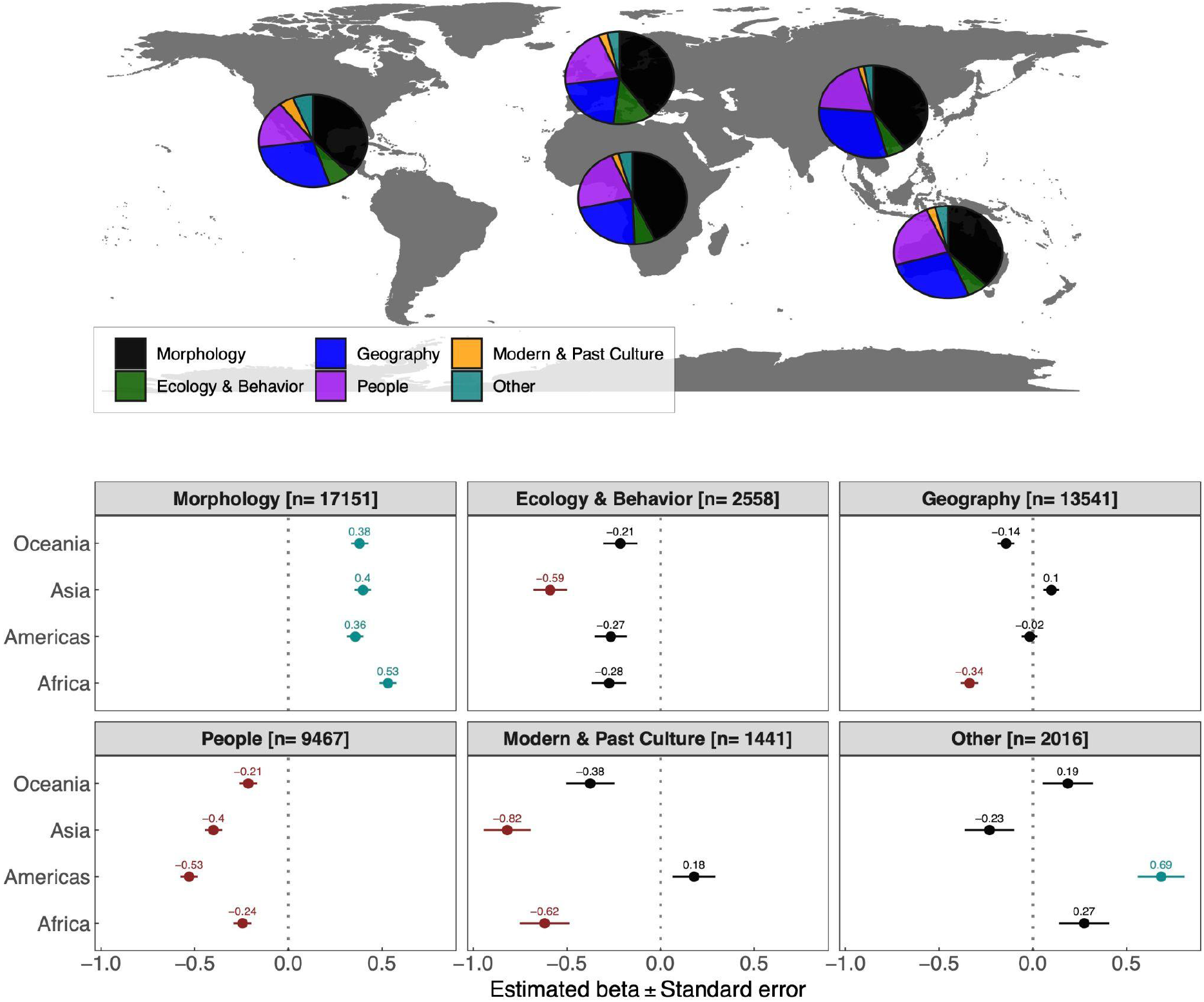
Spatial variations in the way taxonomists assign spiders etymologies. Pie charts display the proportions of etymology types by continent. Estimates (± standard errors) are based on Bernoulli generalized linear models testing for differences in the proportion of different etymology types by continent. Significant differences (p < 0.001) from the baseline (Europe) are highlighted with colors (positive = turquoise; negative = red). Exact model estimates are given in Table S2–S7.

## DISCUSSION

### What’s in a spider name?

Despite gigantic advances in molecular and bioinformatic tools (Tosa *et al*., 2021; Anderson *et al*., 2021), taxonomy remains a conservative science that still mostly relies on morphological methods and the basic grammar rules of the scientific Latin language used in the 18^th^ century. At the same time, it’s a domain of science home to vigorous and even heated debates, for example around the role of taxonomy and taxonomists within Academia (Drew, 2011; Zeppelini *et al*., 2021) and around emerging new approaches to species descriptions [e.g., photography-based taxonomy (Ceríaco & et al., 2016; Zamani *et al*., 2021); sequence-based taxonomy (Sharkey *et al*., 2021; Zamani *et al*., 2022)]. Spider names echo the traditional nature of taxonomy, insofar as most etymologies refer to biological features of spiders and their ecology and distributions. Indeed, the categories of morphology and geography were overwhelmingly represented across the database, both temporally (Figure 1B) and spatially (Figure 2). However, naming practice has changed: names based on morphology, as well as those based on ecology and behavior, have both declined substantially since the late 19^th^ century. Interestingly, the same trend is seen in species etymologies in the plant genus *Aloe* (Figueiredo & Smith, 2010): names from morphology have fallen by about half over the 20^th^ century, while names for geography and people have increased. Figueiredo and Smith speculate that this may reflect either decreasing knowledge of technical Latin or a saturation effect where simple morphological names are increasingly used up. In spiders, at least, we suspect the latter is more likely. However, it is difficult to distinguish this hypothesis from an increasing receptivity to more creative naming, perhaps reflecting broader acceptance that science is, like all human enterprises, culturally situated.

While names based on morphology, geography, and ecology can be seen as deeply rooted in taxonomic tradition, there are some extravagant outliers. For instance, some names combine morphological features and cultural references with unexpected outcomes. Just to cite a few cases, *Heteropoda hippie* Jäger comes from the Hippie movement and refers to the long hairs of the male spider metatarsi (Jäger, 2008); *Eriovixia gryffindori* Ahmed, Khalap & Sumukha, a spider with a hat-shaped abdomen, is a reference to the sorting hat in the Harry Potter books by J.K. Rowling (Ahmed, Khalap, & Sumukha, 2016); and *Heteropoda ninahagen* Jäger is a tribute to the songs of Nina Hagen that, according to the descriptor, are *“[*…*] as unusual as the shape of the retrolateral tibial apophysis”* (Jäger, 2008)—we recommend listening to a few albums to fully appreciate this analogy.

Both in the past and today, a substantial fraction of names were also dedicated to people, with the proportion of such etymologies slowly but steadily increasing between 1757 and 2019. In the past, these names were mostly honoring species collectors or other scientists, especially arachnologists (see Figure S1). In recent years, however, these etymologies have been also used to pay tribute to artists and celebrities or to make statements about politics and the environment. An emblematic case is a study by Agnarsson et al. (2018): in documenting a radiation of *Spintharus* in the Caribbean, they named fifteen species after famous people who stood up for human rights or were committed to nature conservation—Sir David Attenborough (*S. davidattenboroughi* Agnarsson & Van Patten), Barack Obama (*S. barackobamai* Agnarsson & Van Patten), Michelle Obama (*S. michelleobamaae* Agnarsson & Sargeant), and Leonardo Di Caprio (*S. leonardodicaprioi* Van Patten & Agnarsson), among others. Interestingly, in helmith parasites the frequency of eponymous names that honor scientists has not changed, at least over a recent 2-decade window, but the frequency of eponymous names honoring other people has increased (Poulin, McDougall, & Presswell, 2022). Together with our results, which span a much longer period of time, we see this as further evidence of increasing interest in eponymous naming, combined with growing socio-cultural engagement in the coining of names.

In the last two decades, with taxonomy entering the Internet era, there has also been a rise in etymologies dedicated to modern cultures, especially for European and American species. Such etymologies have referred to topics as diverse as the human cultural heritage itself: we found mentions to spiders in fantasy and fantastic literature (e.g., all species described in (Brescovit, Cizauskas, & Mota, 2018), references to cartoons (e.g., *Ianduba dabadu* Magalhaes *et al*. is intended to sound like the famous yell of Fred Flintstone; Magalhaes et al., 2016), tributes to musicians (e.g., *Balmaceda abba* Edwards & Baert, honoring the Swedish pop group ABBA; Edwards & Baert, 2018), allusions to urban legends (e.g., *Antilloides chupacabras* Magalhaes; see detailed explanation in Magalhaes, 2018), and countless other local traditions.

Finally, under the category Other we grouped a miscellany of difficult-to-classify etymologies. Just to cite a few cases, the oonopid spider *Gamasomorpha insomnia* Eichenberger owes its name to several “*sleepless nights*” experienced by the author during discrimination of intraspecific variation and feasible characters (Eichenberger *et al*., 2012); *Hortipes aelurisiepae* Bosselaers & Jocqué is in memory of “… *the cat Siep, which was run over by a truck when this species was being described*” (Bosselaers & Jocqué, 2000); and *Filistatinella howdyall* Magalhaes & Ramírez simply recall the Texan salute “*howdy, y’all*!” (Magalhaes & Ramírez, 2017). Interestingly, 465 of the 2128 etymologies that we classified as Other were “*arbitrary combinations of letters*”. This naming strategy was frequently employed by prolific authors who described thousands of species, including Willis J. Gertsch (1906–1998), Herbert W. Levi (1921–2014), and Norman I. Platnick (1951–2020). While some of these arbitrary names actually have hidden meanings (for example, they were anagrams of the type locality), it was perhaps also a convenient naming strategy when lacking ideas.

### A futile exercise?

One may argue that our classification exercise was of limited and mainly technical interest— especially after knowing that it took us almost two years to go through the full list of spider names! But we do not think so. Besides the narrow-in-scope goal of better understanding naming practices in araneology, we contend that our work can be a benchmark for future studies. More importantly, we believe that our work should foster reflections about the science of taxonomy more generally, and about the way science, and scientists, are situated in a cultural context that influences everything we do.

First, our analysis comes in a historical time when the science of taxonomy is experiencing a profound crisis (Godfray, 2002; Bilton, 2014). Despite gigantic leap forwards in artificial intelligence and molecular technologies, taxonomy—and especially species description—remains a largely conservative discipline, probably one of the most unchanged across the last centuries in terms of background, tools, and rules (but see Ceríaco & et al., 2016; Zamani et al., 2021, 2022; Sharkey et al., 2021). Nowadays, basic taxonomy faces becoming a marginalized discipline, being in constant shortage of personnel and receiving limited research funding, attention, and credit (Godfray, 2002; Engel *et al*., 2021). However, taxonomy is pivotal to any discipline focusing on living organisms and it is fundamental to ensuring reproducibility of biological studies given that species misidentifications or changes in taxonomy may deeply affect conclusions (Vink *et al*., 2012). Although some of the etymologies currently fashionable in modern-day taxonomy carry little or no biological information regarding the species itself, they can help to give visibility to taxonomic and species-discovery research. For example, homages to celebrities—from Johnny Cash (*Aphonopelma johnnycashi* Hamilton) to Angelina Jolie (*Aptostichus angelinajolieae* Bond) to David Bowie (*Heteropoda davidbowie* Jäger and *Spintharus davidbowiei* Agnarsson & Chomitz) to Arnold Schwarzenegger (*Predatoroonops schwarzeneggeri* Brescovit et al.)— often attract huge scientific and media attention (for example, as measured *via* Altmetrics scores). This raises the important question: can this attention be harnessed to spotlight problems surrounding the taxonomy crisis? It would be interesting, but challenging, to ask whether public attention to species naming translates to public or governmental support for species discovery and conservation.

Second, our classification exercise allowed us to discover hidden gems of unconventional and sparkling scientific writing, illustrating how taxonomy’s reputation of being tedious (e.g., Leather & Quicke, 2009) is not entirely deserved. The number of witty etymologies we documented is a reminder that even the writing (and reading) of taxonomic papers can be simultaneously rigorous and enjoyable (Heard, 2014). Also, the recent dramatic increase in non-traditional etymologies is symptomatic of an ongoing transition toward a modern writing style that retreats to some extent from the dense, impersonal, and difficult-to-access orthodox style of scientific writing (Sword, 2012; Greene, 2013; Doubleday & Connell, 2017; Freeling, Doubleday, & Connell, 2019; Mammola, 2020)—otherwise known as “academese” (Pinker, 2015). To us, the creativity of taxonomists in devising new species names is inspirational and can foster reflection about the way we communicate science.

Third, our database can be a starting point to explore important topics related to academic life, especially the timely debate around inequalities within the scientific community. An illustrative example comes from a recent work by Trisos et al. (2021). They mapped Latin names of birds globally, showing that most of the species occurring outside Europe in formerly colonized countries are named after European surnames rather than indigenous ones. In our case, a subset of spider etymologies dedicated to people could be used to confirm the generality of this pattern beyond birds. Also, one could test whether there is a gender bias in the way taxonomists assign species names and if the ratio male:female in names is changing through time. We believe the latter would be a timely endeavor given that reflections around gender issues are gaining ground within the arachnological community (check, for example, the multiple initiatives by the recently founded group SWA–Support Women in Arachnology; @SwaWomen). We leave the answers to these and other questions for future work while welcoming anyone to explore and re-use the database uploaded alongside this submission. For now, we simply highlight two recent examples: *Grymeus dharmapriyai* Ranasinghe & Benjamin and *Tmarus manojkaushalyai* Ileperuma Arachchi & Benjamin—each named by a young Sri Lankan woman, as part of her taxonomic training, for her husband (Ranasinghe & Benjamin, 2018; Ileperuma Arachchi & Benjamin, 2019). Here, taxonomy and creativity come together to express perhaps the best of human emotions: love. Science and scientists will always be embedded in human culture, and because they allow such free expression, scientific names can be a way to shine a light on this connection.

## SUPPLEMENTARY MATERIALS

Table S1–S7

Figure S1–S2

## ACKNOWLEDGMENTS

We are grateful to the World Spider Catalog’s team (https://wsc.nmbe.ch/boards) for their immense contribution to spider taxonomy. Special thanks to people who helped us with inferring etymologies in specific languages, namely Malin Teräväinen (Swedish) and Svetlana Pétillon (Russian).

## DATA AVAILABILITY STATEMENT

R codes to generate the analyses and plots is available in GitHub (https://github.com/StefanoMammola/Spider_Etymologies_Analysis). The database supporting the study is available in Figshare (https://doi.org/10.6084/m9.figshare.19126658). Link will start working upon acceptance in an official journal.

## AUTHOR CONTRIBUTION

JP conceived the idea. JP, DF, and SM designed methodology. All authors collected data. SM analyzed data and wrote the first draft. All authors contributed to the writing with suggestions, ideas, and additions to the text.

## FIGURES AND BOXES

### Box 1.

**Spider etymologies by numbers and some Spider World Records**

The number of characters in spider names is roughly normally distributed, considering either the sum of letters in the genus and species (Figure box A) or the species epithet alone (Figure box B). The average number of characters in the species epithet hasn’t changed substantially between 1757 and today (Figure box C; note that the variability in the average prior 1850 is largely attributable to the lower sample size). When considering the sum of letters in the genus and species, the record for the longest (spider) scientific name goes to the recently described theraphosid spider *Chilobrachys jonitriantisvansickleae* Nanayakkara, Sumanapala & Kirk (35 characters), dedicated to Joni Triantis Van Sickle (Nanayakkara, Sumanapala, & Kirk, 2019). This newly described species beat the previous records of 33 characters (Mammola *et al*., 2017), which was held by *Dipoena santaritadopassaquatrensis* Rodrigues (Theridiidae) and *Phormingochilus pennellhewlettorum* Smith & Jacobi (Theraphosidae). When only taking into account the species epithet, however, *D. santaritadopassaquatrensis* still yields the record for the longest name (26 characters). The etymology is derived from the type locality, Santa Rita do Passa Quatro in the state of São Paulo, Brasil (Rodrigues, 2013). Conversely, the shortest etymology (Genus + species) is six letters in *Gea eff* Levi (Araneidae) (Mammola *et al*., 2017). According to the original description, *eff* is simply an “*arbitrary combination of letters*” (Levi, 1983). Several two-letter epithets have also been attributed to various species (*ab, an, ef, fo, la, kh, mi, no, oz, wa, wu, yi*, and *zu*). Interestingly, some of these short epithets are used intentionally to refer to the small size of the species. For example, *Selenops ab* Logunov & Jäger (Selenopidae) and *Selenops ef* Jäger (Selenopidae) are arbitrary combinations of letters reflecting the small size of the species by using the smallest allowable number of letters to form a specific name (Lagunov & Jäger, 2015; Jäger, 2019). It is worth noting that *ab* is currently also the first spider epithet alphabetically. This, until someone names a species *aa*: the challenge is open!

**Figure Box.**
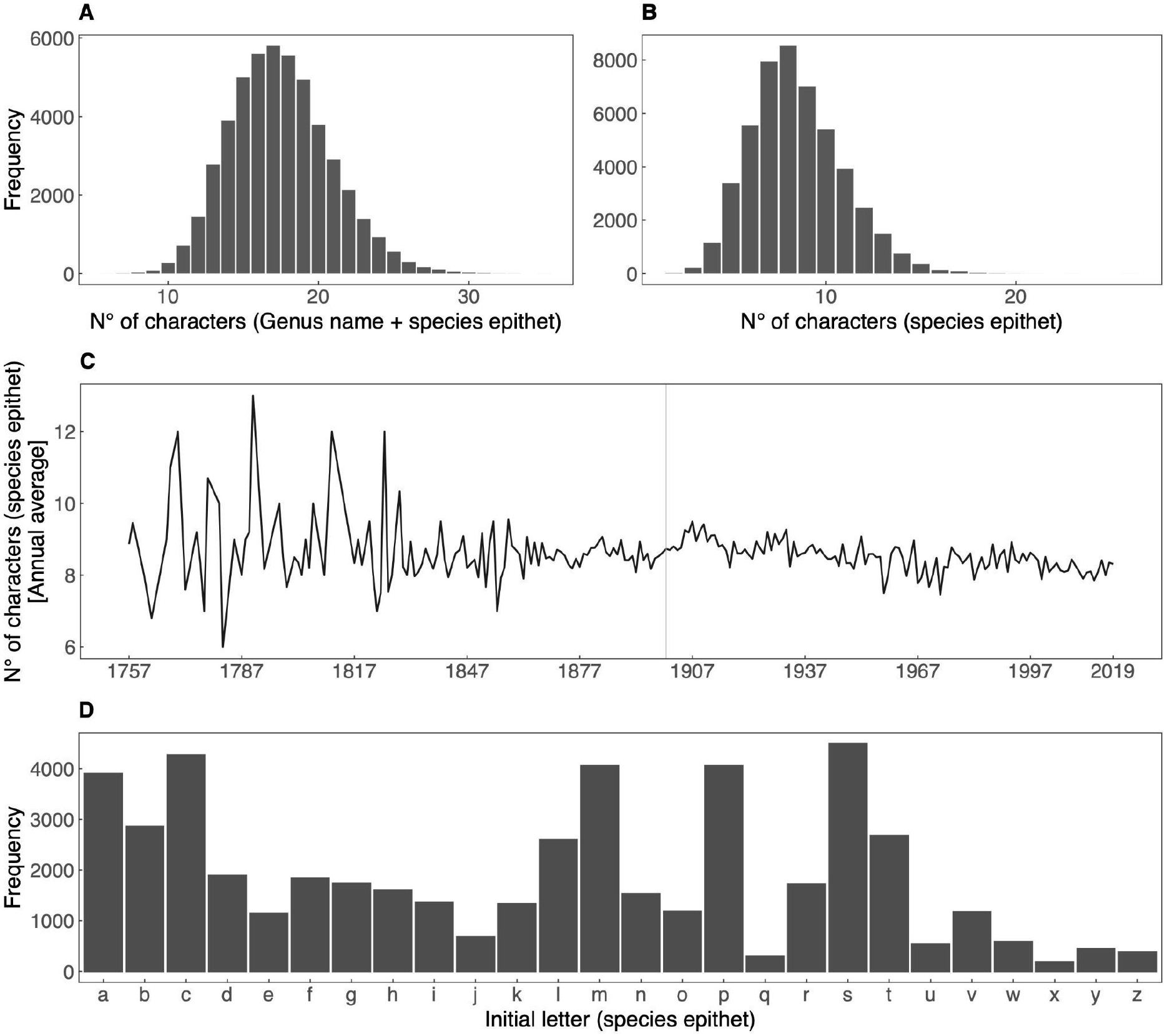
**A)** Frequency of etymologies by the number of letters, summing genus plus species. **B**) Frequency of etymologies by the number of letters in the species epithet. **C**) Variation in the length of species epithets (annual average) over time (1757–2019). **D**) Frequency of etymologies by the initial letter of the species epithet.

