## SUPPLEMENTARY MATERIAL for "Taxonomic practice, creativity, and fashion: What’s in a spider name?"

**Supplementary materials include:**

Table S1–S7

Figure S1–S2

**Table S1. Estimated model parameters for the generalized additive model testing the effect of a smooth function of year of publication in interaction with the etymology category on the proportion of etymologies.** Smoothing functions are shown in Figure 2C. edf = estimated degrees of freedom.

| Component | Term | Estimate β | SE | t-value |
|---|---|---|---|---|
| Parametric coefficients | (Intercept) | -0.750 | 0.026 | -29.317 |
| | Type [Ecology & Behavior] | -1.708 | 0.052 | -32.790 |
| | Type [Geography] | -1.245 | 0.087 | -14.252 |
| | Type [People] | -1.211 | 0.063 | -19.356 |
| | Type [Modern & Past culture] | -3.602 | 0.143 | -25.256 |
| | Type [Other] | -2.507 | 0.102 | -24.629 |
| **Component** | **Term** | **edf** | **Ref. df** | **F-value** |
| Smooth terms | s(Year) | 1.002 | 1.003 | 9.204 |
| | s(Year) * Type [Morphology] | 5.413 | 6.455 | 12.340 |
| | s(Year) * Type [Ecology & Behavior] | 5.125 | 6.177 | 3.879 |
| | s(Year) * Type [Geography] | 8.199 | 8.695 | 15.401 |
| | s(Year) * Type [People] | 5.630 | 6.608 | 13.869 |
| | s(Year) * Type [Modern & Past culture] | 3.421 | 4.234 | 20.254 |
| | s(Year) * Type [Other] | 5.663 | 6.667 | 8.849 |

**Table S2. Estimated parameters for the Bernoulli generalized linear model testing the effect of the factor continent on the use of the etymology type Morphology.** The level 'Europe' is the baseline for the factor continent. SE = Standard Error.

| | **Estimate β** | **SE** | **z value** | **Pr(>\|z\|)** |
|---|---|---|---|---|
| (Intercept) | -1.112 | 0.042 | -26.464 | - |
| Continent [Africa] | 0.532 | 0.046 | 11.637 | <0.0001 |
| Continent [Americas] | 0.357 | 0.044 | 8.098 | <0.0001 |
| Continent [Asia] | 0.398 | 0.044 | 8.954 | <0.0001 |
| Continent [Oceania] | 0.380 | 0.046 | 8.206 | <0.0001 |

**Table S3. Estimated parameters for the Bernoulli generalized linear model testing the effect of the factor continent on the use of the etymology type Ecology & Behavior.** The level 'Europe' is the baseline for the factor continent. SE = Standard Error.

| | **Estimated β** | **SE** | **z value** | **Pr(>\|z\|)** |
|---|---|---|---|---|
| (Intercept) | -2.504 | 0.079 | -31.664 | - |
| Continent [Africa] | -0.275 | 0.092 | -2.991 | 0.0028 |
| Continent [Americas] | -0.266 | 0.086 | -3.105 | 0.0019 |
| Continent [Asia] | -0.589 | 0.090 | -6.579 | <0.0001 |
| Continent [Oceania] | -0.214 | 0.092 | -2.326 | 0.0200 |

**Table S4. Estimated parameters for the Bernoulli generalized linear model testing the effect of the factor continent on the use of the etymology type Geography.** The level 'Europe' is the baseline for the factor continent. SE = Standard Error.

| | **Estimated β** | **SE** | **z value** | **Pr(>\|z\|)** |
|---|---|---|---|---|
| (Intercept) | -0.968 | 0.040 | -24.421 | - |
| Continent [Africa] | -0.338 | 0.046 | -7.307 | <0.0001 |
| Continent [Americas] | -0.017 | 0.042 | -0.404 | 0.6861 |
| Continent [Asia] | 0.099 | 0.042 | 2.331 | 0.0198 |
| Continent [Oceania] | -0.143 | 0.046 | -3.141 | 0.0017 |

**Table S5. Estimated parameters for the Bernoulli generalized linear model testing the effect of the factor continent on the use of the etymology type People.** The level 'Europe' is the baseline for the factor continent. SE = Standard Error.

| | **Estimated β** | **SE** | **z value** | **Pr(>\|z\|)** |
|---|---|---|---|---|
| (Intercept) | -1.072 | 0.041 | -25.923 | - |
| Continent [Africa] | -0.245 | 0.048 | -5.127 | <0.0001 |
| Continent [Americas] | -0.530 | 0.045 | -11.657 | <0.0001 |
| Continent [Asia] | -0.400 | 0.046 | -8.736 | <0.0001 |
| Continent [Oceania] | -0.214 | 0.048 | -4.468 | <0.0001 |

**Table S6. Estimated parameters for the Bernoulli generalized linear model testing the effect of the factor continent on the use of the etymology type Modern & Past Culture.** The level 'Europe' is the baseline for the factor continent. SE = Standard Error.

| | **Estimated β** | **SE** | **z value** | **Pr(>\|z\|)** |
|---|---|---|---|---|
| (Intercept) | -3.168 | 0.109 | -29.033 | - |
| Continent [Africa] | -0.618 | 0.134 | -4.627 | <0.0001 |
| Continent [Americas] | 0.178 | 0.115 | 1.550 | 0.1211 |
| Continent [Asia] | -0.818 | 0.127 | -6.435 | <0.0001 |
| Continent [Oceania] | -0.375 | 0.130 | -2.885 | 0.0039 |

**Table S7.** **Estimated parameters for the Bernoulli generalized linear model testing the effect of the factor continent on the use of the etymology type Others.** The level 'Europe' is the baseline for the factor continent. SE = Standard Error.

| | **Estimated β** | **SE** | **z value** | **Pr(>\|z\|)** |
|---|---|---|---|---|
| (Intercept) | -3.383 | 0.121 | -27.898 | - |
| Continent [Africa] | 0.274 | 0.133 | 2.055 | 0.0399 |
| Continent [Americas] | 0.685 | 0.125 | 5.467 | <0.0001 |
| Continent [Asia] | -0.231 | 0.133 | -1.740 | 0.0819 |
| Continent [Oceania] | 0.186 | 0.135 | 1.378 | 0.1683 |

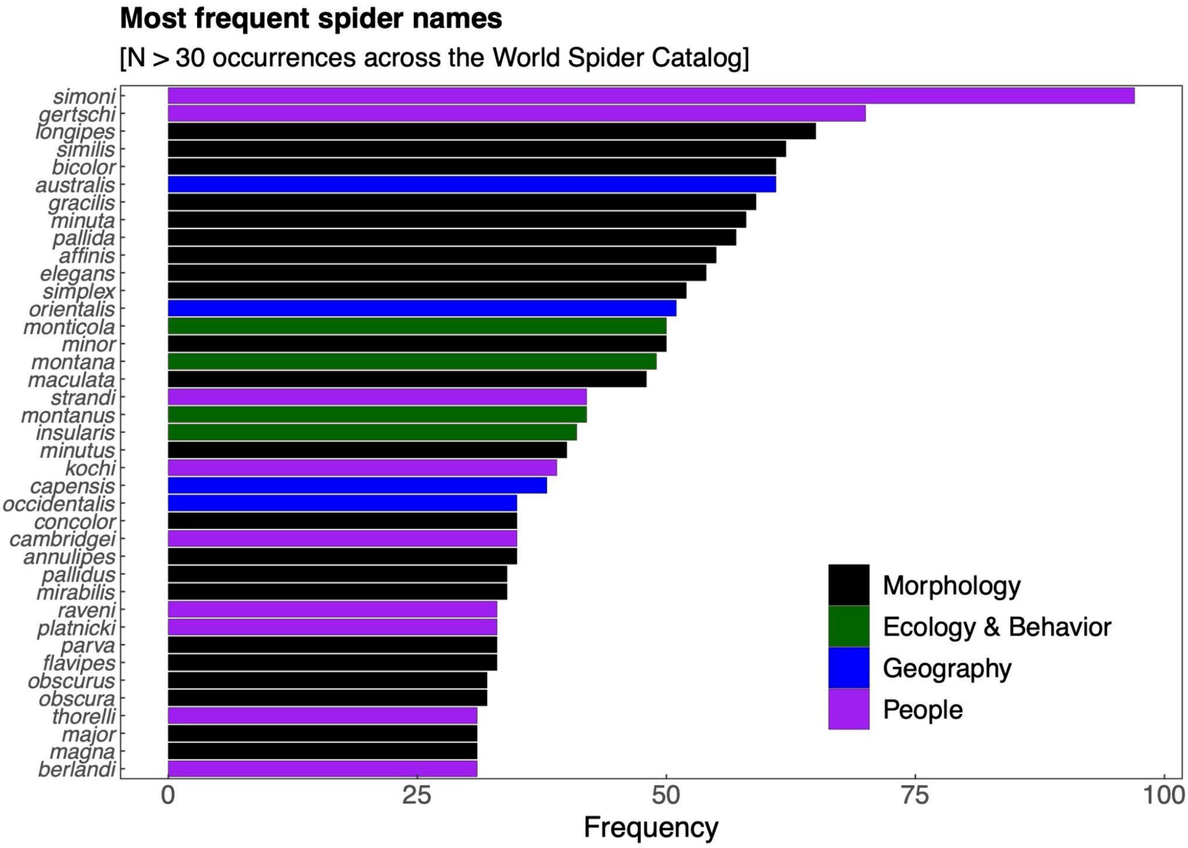


**Figure S1. Most frequent spider epithets.** Only epithets shared by more than 30 species across the database are shown. Color coding refers to the most frequent use of each name (e.g., *similis* is most often used in a morphological sense, but in a few instances it was also used as a reference to the similarity in the geographic distribution of one species to another).

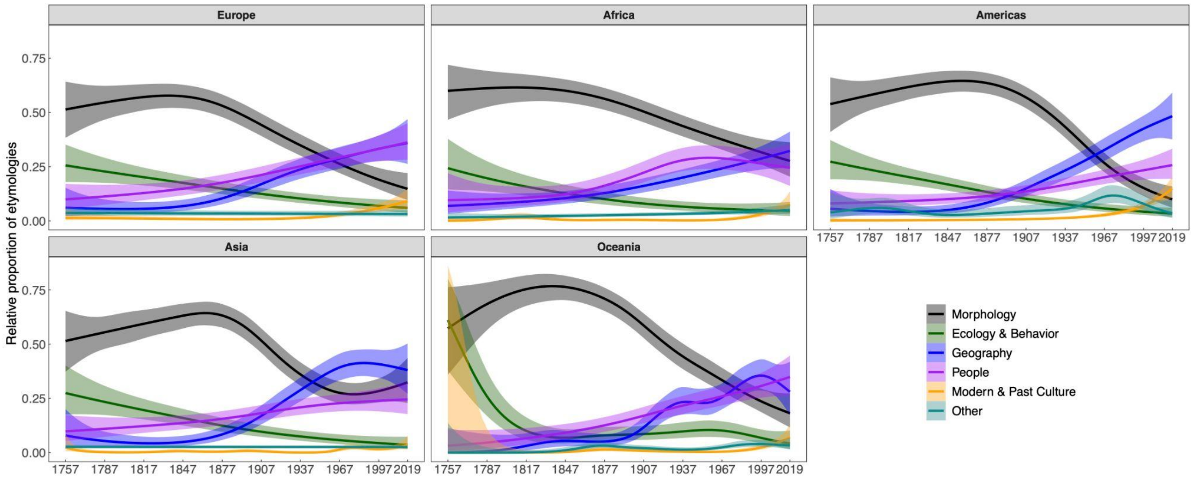


**Figure S2.** **Temporal trends in the way taxonomists assign etymology by region.** Colored lines are predicted values and shaded surfaces represent 95% confidence intervals.
